# Benchmarking algorithms for spatially variable gene identification in spatial transcriptomics

**DOI:** 10.1101/2024.07.04.602147

**Authors:** Xuanwei Chen, Qinghua Ran, Junjie Tang, Zihao Chen, Siyuan Huang, Xingjie Shi, Ruibin Xi

**Author notes:** Corresponding author: Dr. Ruibin Xi, No.5 Yiheyuan Road Haidian District, Beijing, 100871, China, Dr. Xingjie Shi, No.3663 North Zhongshan Road, Putuo District, Shanghai, 200062, China.

## Abstract

The rapid development of spatial transcriptomics has underscored the importance of identifying spatially variable genes. As a fundamental task in spatial transcriptomic data analysis, spatially variable gene identification has been extensively studied. However, the lack of comprehensive benchmark makes it difficult to validate the effectiveness of various algorithms scattered across a large number of studies with real-world datasets. In response, this article proposes a benchmark framework to evaluate algorithms for identifying spatially variable genes through the analysis of synthesized and real-world datasets, aiming to identify the best algorithms and their corresponding application scenarios. This framework can assist medical and life scientists in selecting suitable algorithms for their research, while also aid bioinformatics scientists in developing more powerful and efficient computational methods in spatial transcriptomic research.

## Introduction

In recent years, the rapid development of spatial transcriptomics has enabled simultaneous measurements of the molecular expression of mRNAs and their corresponding spatial information within a sample [1, 2]. The studies of spatial transcriptomic data have enhanced our understanding of tissue morphology, tissue microenvironment and inter-cellular communication mechanisms [3–7]. Recognized as the “method of the year” by Nature in 2020 [8], the field of spatial transcriptomics has experienced an explosion of studies, leading to the generation of increasingly complex datasets with varying resolutions. Additionally, large-scale public data sources, such as whole mouse brain atlas [9], Spatial Omics DataBase (SODB) [10], and Spatial Tumor MicroEnvironment (SpatialTME) database [11], have become available. Many computational methods have been developed for spatial transcriptomic data, leading to significant research advancements in this field [12–14].

One of the fundamental and crucial analysis tasks in spatial transcriptomics is identifying spatially variable genes (SVGs), which are genes exhibiting spatial patterns of expression variations [15]. In contrast to highly variable genes (HVGs) in single-cell RNA sequencing (scRNA-seq) data, SVGs are more concerned with the dependence of gene expression variations on spatial locations. The identification of SVGs provides insights into essential biological functions and mechanisms related to spatial tissue morphology, differentiation and inter-cellular communication [16–18]. Furthermore, identification of SVGs can enhance downstream analysis tasks, such as spatial domain detection [19], spatially-aware dimension reduction [20] and trajectory inference [21].

To enable accurate detection of SVGs with diverse spatial patterns [15, 22, 23], and address the challenges of large-scale spatial transcriptomic data [24–26], many SVG methods have been developed in recent years. According to the ways in which spatial information is captured, these methods can be roughly classified into four categories: (1) Kernel-based methods, which utilize kernel functions to organize spatial distance and employ random field models to capture spatial dependencies of expression. SpatialDE [15] employs a Gaussian process regression model to test spatial variance. SPARK [23] uses a quasi-Poisson generalized linear spatial model with a logarithmic link function and utilizes 10 candidate variance kernels to enhance stability. Based on a Gaussian process regression model, GPcounts [27] and BOOST-GP [28] further model counts using negative binomial (NB) and zero-inflated negative binomial (ZINB) distributions, respectively. SOMDE [26] adopts a self-organizing map to merge adjacent spatial localization on the basis of SpatialDE, enhancing computational efficiency for large-scale data. (2) Marked point process methods with spatial locations serving as points and gene expression as marks. Largely speaking, trendsceek [22] models the joint probability distribution of the expression levels and spatial locations, and uses random permutation to construct null model for testing. scGCO [29] utilizes hidden Markov random field (HMRF) and graph-cutting to identify SVGs. (3) Correlation-based methods that assess correlation between gene expression values and spatial coordinates. SPARK-X [30] defines a type of correlation between gene expression values and spatial coordinates, involving multiple transformations of spatial coordinates to capture various types of expression patterns. BinSpect [31] uses Fisher’s exact test to examine whether binarized expression exhibits any spatial auto-correlation patterns. MERINGUE [32] uses Moran’s index and Local Moran’s index (LISA) to capture global and local spatial autocorrelation. (4) Miscellany methods. BOOST-MI [33] applies a modified Ising model to model binarized expression profiles, and uses energy interaction parameters to indicate whether there are spatial differences in gene expression. sepal [34] proposes to distinguish SVGs from non-SVGs by diffusion equations and assumes that spatial variability is directly proportional to diffusion time. See Methods for more details.

Given the diversity of the available SVG methods, it is crucial to comprehensively assess their performances across various aspects, such as their detection accuracy, statistical validity, stability, and scalability. Moreover, it is important to provide a detailed user guide in selecting SVG identification methods, tailored to specific characteristics of spatial transcriptomic data and analytical requirements. Although there are a few benchmarking studies [35, 36], their evaluations are constrained by the use of a limited small number of real-world datasets (n<25), an insufficient range of evaluation criteria, and the absence of an informative user guide that addresses the specific analytical needs and data scenarios encountered in practice. Therefore, there is an urgent need for a more comprehensive comparison of SVG methods.

We develop a comprehensive benchmark framework that evaluates 15 SVG identification methods (Methods) using 30 synthesized and 74 curated real datasets from various technologies (Fig. 1, Table S1). This framework evaluates the SVG methods in terms of SVG detection accuracy, statistical validity, accuracy of downstream clustering, stability and scalability. Furthermore, we offer a context-specific user guide that recommends methods based on specific analysis requirements and data characteristics.

**Fig. 1.**
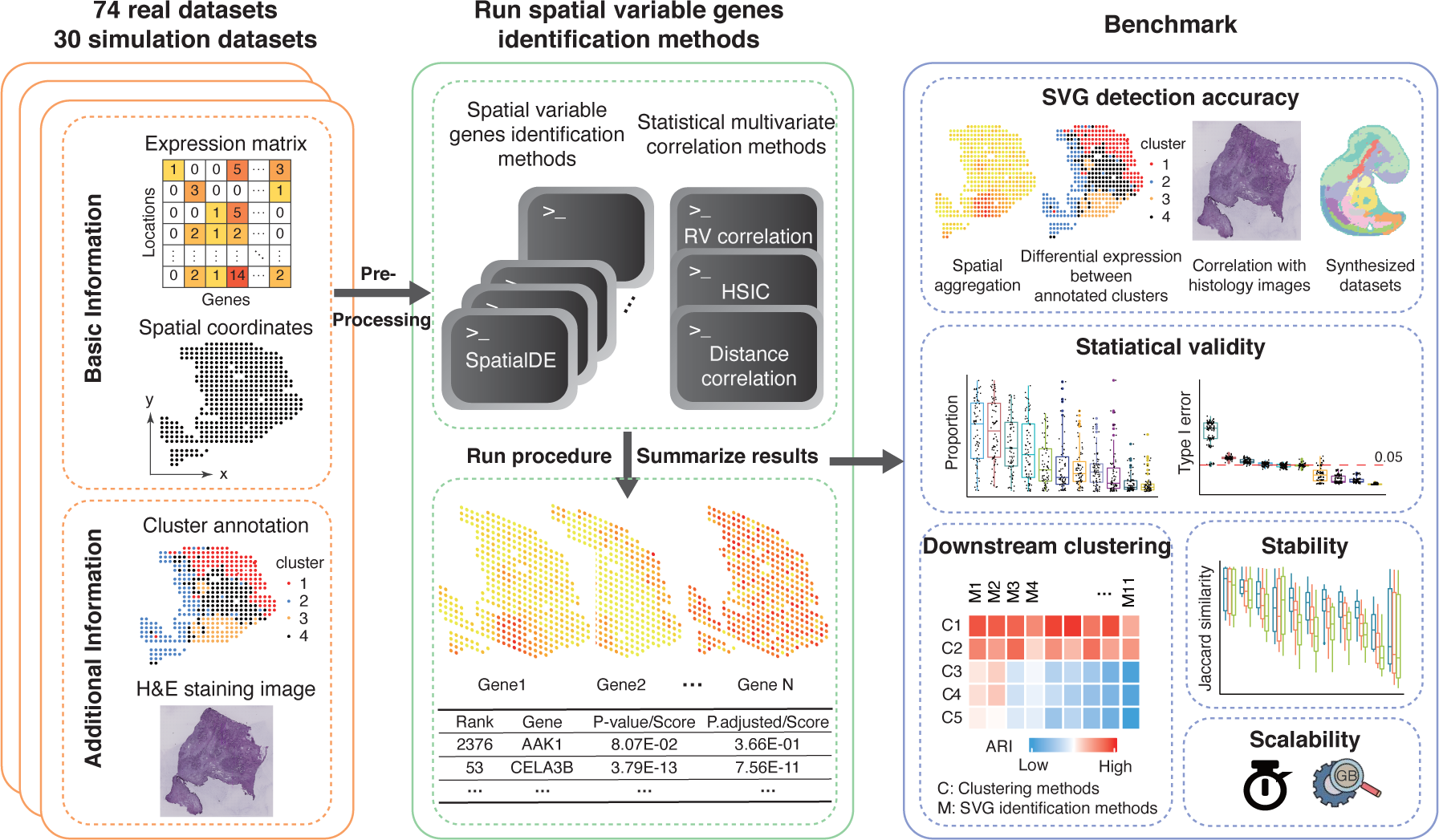
An overview of the benchmark framework. We first pre-processed each dataset through a uniform pipeline, and then applied 15 candidate SVG identification algorithms to both real-world datasets from curated literature and synthetic datasets generated by SRTsim based on actual experiment data. For each method, crucial results were extracted and summarized, including the SVG ranks, corresponding p-values/scores and adjusted p-values/normalized scores. we evaluated algorithms in terms of SVG detection accuracy, statistical validity, accuracy of downstream clustering, stability and scalability (time and memory).

## Results

### Overview of Benchmark Framework

To comprehensively assess the performance of various SVG identification methods, we collected 74 real-world datasets encompassing a wide range of protocols, tissues, data sizes and resolutions (Fig. S1, Table S1). We also collected associated external features for these datasets, such as cluster annotations, histology images, and data from adjacent slices, to aid in developing comprehensive evaluation criteria (Methods). In addition, we generated 30 synthesized datasets to simulate spatial transcriptomic data from three representative technologies, 10x Visium [37], Stereo-seq [38], and Slide-seqV2 [39]. To obtain gold standard SVGs on the synthesized data, we first employed the estimated parameters from the simulator SRTsim [40] to categorize genes into SVG sets, non-SVG sets, and ambiguous gene sets. We then excluded the ambiguous gene sets, which include genes that fall on the boundary of SVGs or non-SVGs, to avoid evaluation bias. Using the simulator SRTsim, we generated 10 synthesized datasets for each reference dataset, ensuring that the synthesized data matched the number of spatial locations, resolutions, dropout rates and other relevant properties of the corresponding reference dataset (Methods; Fig. S2). These synthesized and real-world datasets served as a representative sample of the complexities typically encountered in real-world spatial transcriptomic experiments.

We conducted an extensive literature review on SVG identification algorithms. After excluding methods [41–43] that combine two distinct tasks - spatial region identification and spatial variable gene detection - or lacking user manuals, we obtained 15 candidate methods for benchmarking. The 15 methods included 12 algorithms specially designed for SVG detection, and 3 additional general-purpose multivariate-correlation methods: RV-coefficient [44], distance correlation (dCor) coefficient [45] and Hilbert Schmidt Independent Criterion (HSIC) [46]. We adapted the general-purpose methods for SVG detection by directly examining the correlation between gene expression and spatial coordinates (Methods; Table S2). We evaluated each method based on five core aspects (Fig. 1) : (1) SVG detection accuracy, obtained by comparing the identified SVGs with silver standard SVGs in real datasets or with gold standard SVGs in synthetic datasets; (2) statistical validity, including the SVG proportions and the type I error rates; (3) accuracy of downstream clustering compared to available annotations; (4) stability, including the reproducibility of identified SVGs between adjacent slices, and the robustness of identified SVGs against spatial location swapping in the datasets; and (5) scalability in terms of time cost and computational memory. In the benchmarking, we imposed a 24-hour time limit. Most methods successfully completed the SVG analysis across the majority of the datasets (Fig. S3). However, GPcounts [27], trendsceek [22], BOOST-MI [33] and BOOST-GP [28] only succeeded in less than 10% of the datasets, and therefore were excluded from further performance comparisons. Consequently, a total of 11 methods were included in our benchmark.

### Accuracy

We evaluated the accuracy of SVG identification methods using both real datasets and synthetic datasets (Methods). In the real data benchmark, we constructed four sets of silver standard SVGs by investigating whether the genes exhibited significant spatial aggregation in terms of Moran’s index, differential expression between known annotated clusters as determined by the Wilcoxon test or the likelihood ratio test of the NB regression model, or correlation with histology images (examples: Fig. S4). Given the silver standard SVGs in real datasets and true SVGs in synthetic datasets, we ranked genes by adjusted p-values or scores provided by each method and computed several widely recognized accuracy metrics (see Methods for details): area under the precision-recall curve (AUPR), area under the receiver operating characteristic curve (AUROC), and early precision ratio (EP). EP is calculated as the fraction of true positives in the top-K identified SVGs, where K is the number of the sliver SVGs. Higher values in these three accuracy metrics indicate better performance of the method.

Overall, BinSpect, SPARK, SpatialDE, dCor, and SPARK-X were the top-performing methods across multiple metrics on both real-world and synthetic datasets (Fig. 2, Fig. S5-6). With respect to the spatial-aggregation silver standards, BinSpect and MERINGUE exhibited the best performance. MERINGUE’s good performance was expected, since it identified SVGs based on the global and local Moran’s index, which was very similar to our way of constructing the spatial-aggregation silver standards. For the silver standards constructed by differential expression analysis between known annotation clusters, BinSpect, SPARK-X, dCor, SPARK, and HSIC generally performed well. Since the Wilcoxon test is a distribution-free statistical test and the likelihood ratio test of NB regression is a parametric test, the silver standards constructed using the Wilcoxon test tended to be more conservative (Fig. S7). As a result, the distribution-free methods BinSpect, HSIC, and dCor performed better for the silver standards constructed by the Wilcoxon test, while SPARK-X, SPARK and sepal outperformed other methods under the NB-regression silver standards. In terms of the silver standards constructed by correlating gene expression with histological staining patterns, BinSpect, SPARK, HSIC and SpatialDE were the best-performing methods. The evaluation results using the synthetic data were largely consistent with the results based on real datasets, with BinSpect being the best-performing method (Fig. 2, Fig. S5). Additionally, when analyzing the “Tissue3” synthetic data simulated from Slide-seqV2 [39], only four methods (BinSpect, SPARK-X, SOMDE, and sepal) were able to successfully complete the analyses (Tissue3 in Fig. 2). The “Tissue3” data boasted an impressive 51,200 spatial locations, highlighting the computational scalability challenges faced by other methods in handling large datasets (Fig. S5). The scalability will be further explored in the ‘Scalability’ section.

**Fig. 2.**
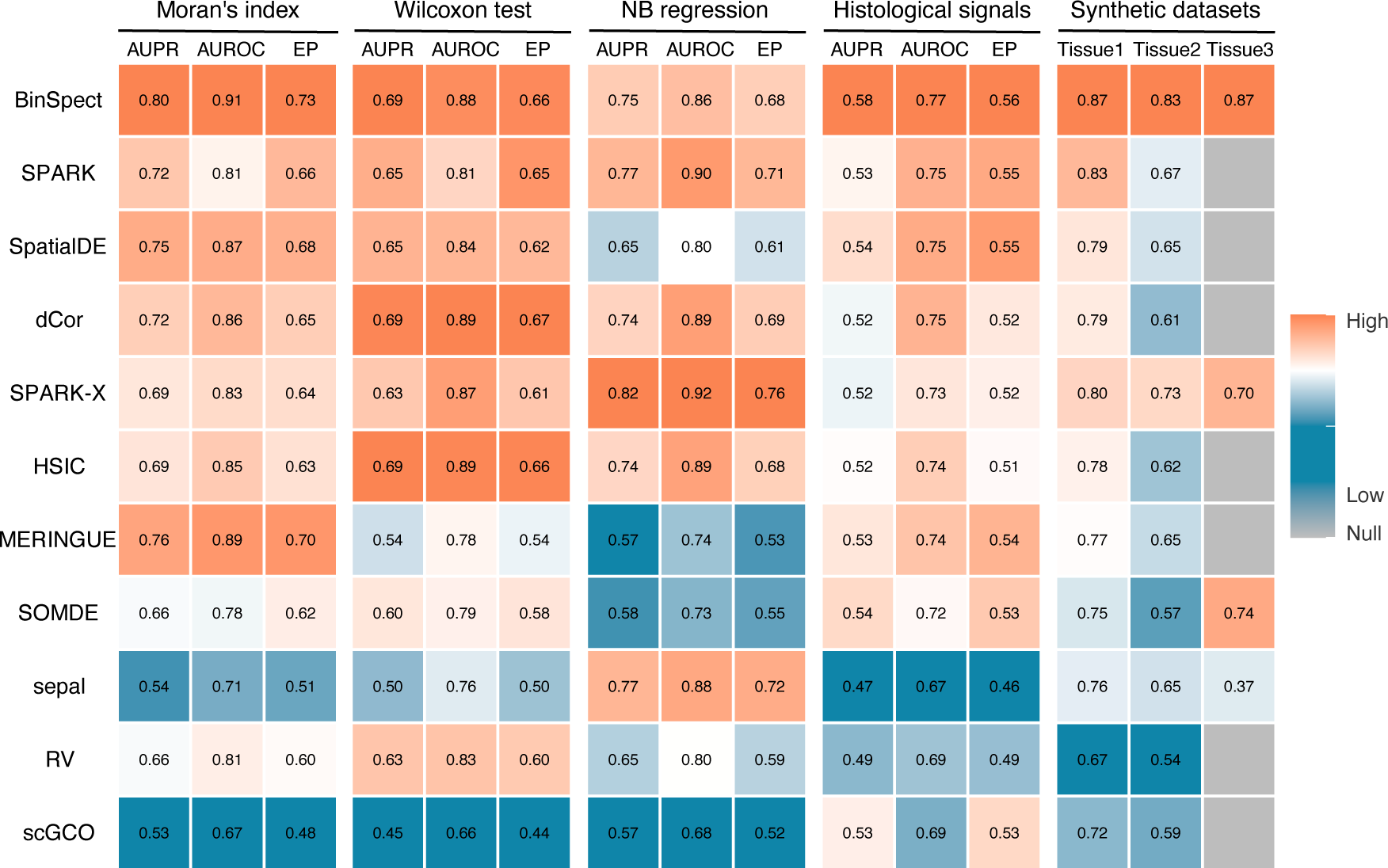
Accuracy of SVG detection. The first twelve columns display the results of accuracy analysis based on the silver standard SVG sets constructed using the Moran’s index, the Wilcoxon test, the NB regression, and the correlation with histological images, respectively. Among them, the accuracy analysis of each silver standard includes the mean values of three accuracy metrics (AUPR, AUROC, and EP), with larger accuracy metrics indicating better performance. The last three columns show the accuracy obtained using synthetic data simulated from different tissues (Tissue1: mouse brain anterior from 10x Visium [37], Tissue2: mouse embryo from Stereo-seq [38], Tissue3: mouse hippocampus from Slide-seqV2 [39]). The colors in the heatmap represent column-wise normalized values of the numbers in the corresponding cells (scaled between 0 and 1). Cells in grey indicate that the corresponding analyses failed to complete. Algorithms (rows) are arranged in descending order of the average rankings.

To examine the potential influence of spatial resolution or the diameter of individual locations, we categorized the real datasets into three groups based on resolution: high-resolution (<20μm), moderate-resolution (20-50μm), and low-resolution (>50μm). We then compared the performance of SVG identification methods within each resolution group by evaluating their Moran’s indices (Fig. S8). Most methods maintained consistent performance levels in identifying SVGs across high-, moderate-, and low-resolution datasets, with BinSpect showing the best performance. In contrast, MERINGUE and SpatialDE demonstrated lower Moran’s indices in the high-resolution datasets, which may be due to the high sparsity of these datasets. SOMDE, on the other hand, achieved better performance in medium and high-resolution datasets, which may be attributed to the effect of merging adjacent locations by the self-organizing map used in this method.

### Statistical validity

We compared the proportions of SVGs called by different methods at a nominal false discovery rate (FDR) cutoff of 0.05 in the real data experiments (Methods). Most methods reported about 5% to 25% of genes as SVGs (Fig. 3A, Table S3). Notably, HSIC, dCor, BinSpect, and SPARK-X reported the highest proportion of genes as SVGs, with an average of over 35% of genes identified as SVGs. These four methods all detected SVGs by investigating correlations between gene expression and spatial locations, indicating that the correlation-based methods may have higher FDRs than other methods. Among the methods specifically developed for SVG detection, BinSpect identified the largest proportion of genes as SVGs (about 42%), while scGCO reported the smallest proportion of SVGs (about 6%).

**Fig. 3.**
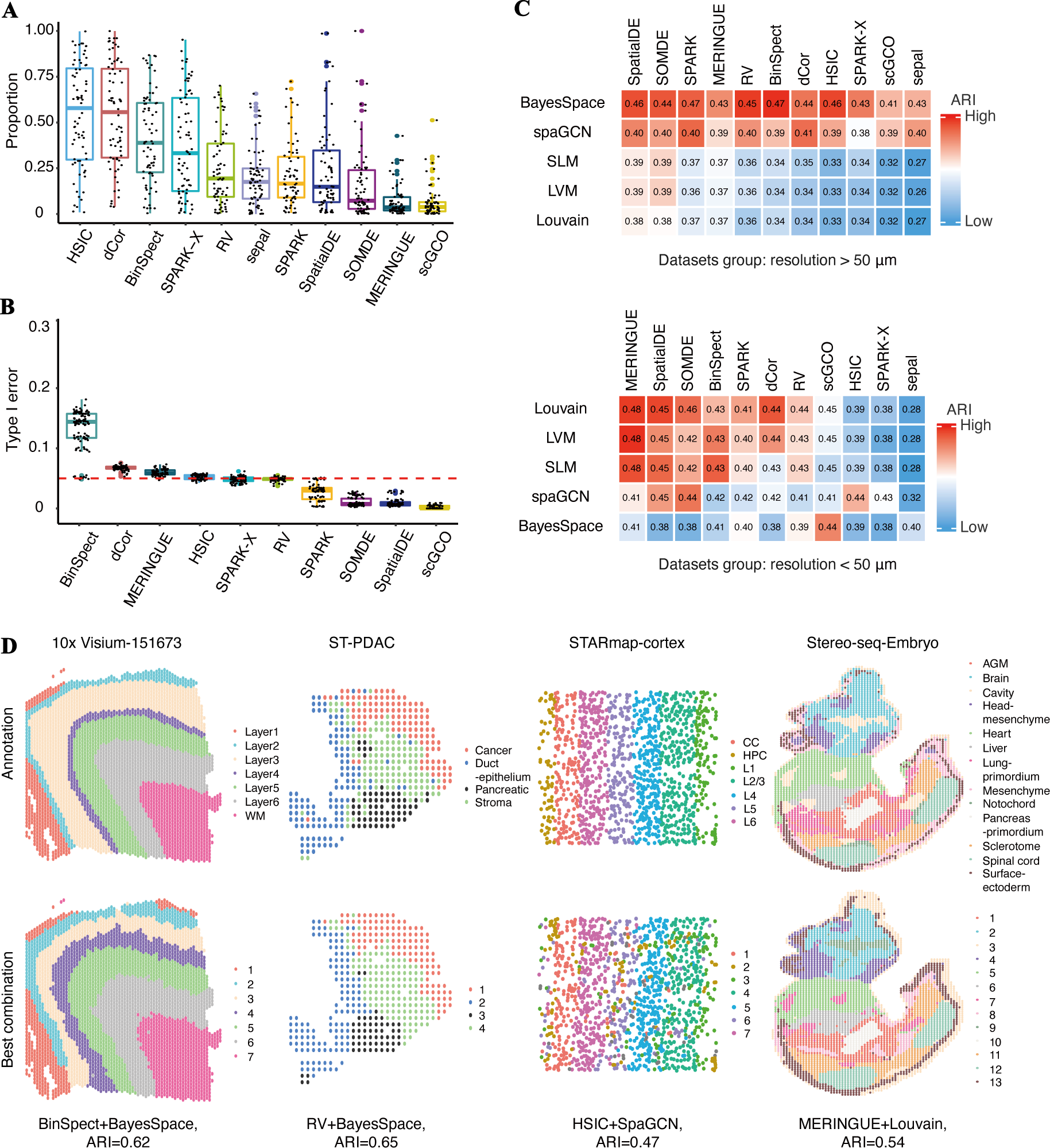
Statistical validity and clustering accuracy. **A**, Box plots of the proportions of SVGs identified by different methods. **B**, Box plots of the type I errors on pseudo-datasets at a nominal p-value of 0.05 (red line). **C**, The heatmap of average ARIs for combinations of clustering methods (rows) and SVG identification methods (columns) for low-resolution (top panel) and high-resolution (bottom panel) real datasets. Top 2000 SVGs were used for the clustering analysis. The colors in the heatmap represent the overall ranking of the ARI for each combination, and the values in the cells are the average ARIs. **D**, Representative clustering results for four different technologies, with each color representing a cluster. Top panel: the known cluster annotations. Bottom panel: the cluster results given by the best-performing combination of clustering and SVG methods.

We then used random permutation to evaluate the type I error rates of the SVG methods (the proportion of non-SVGs incorrectly identified as SVGs). Specifically, we generated pseudo datasets by randomly shuffling and rearranging spatial locations of gene expression measurements across 11 real datasets spanning various technologies (Table S1, Sheet2: ref. data for robustness). Then, for each method, we calculated the proportion of genes reported as SVGs at a nominal p-value cutoff of 0.05, which served as an empirical estimate of the type I error rate (Methods). Other than BinSpect, dCor, MERINGUE, and HSIC, most methods can effectively control the type I errors (Fig. 3B). Among the four methods that cannot control the type I errors, BinSpect showed significant type I errors exceeding the nominal threshold, and its type I errors consistently exceeded 0.1 across datasets, indicating that BinSpect may have introduced an excessive number of false positives during SVG identification (Fig. S9).

### Performance on downstream spatial clustering

Subsequent to SVG identification, a common and crucial analytical task in spatial transcriptomics is to leverage these SVGs for delineating spatially distinct regions or domains within the tissue [2]. We extracted the top 2,000 SVGs identified by SVG methods, and then employed five popular clustering methods to obtain spatial clusters. The five clustering methods included two types of clustering approaches commonly used in spatial transcriptomic studies, state-of-the-art scRNA-seq clustering methods (Louvain algorithm [47], LVM [48], SLM [48]), and spatially-aware clustering methods tailored to spatial transcriptomics (BayesSpace [42], spaGCN [41]). In order to compare the performance of combinations (SVG identification methods + clustering methods) in spatial clustering tasks, we calculated the Adjusted Rand index [49] (ARI) between the obtained clusters and the known cluster annotations (Fig. 3C and Fig. S10-12).

We found that the spatial resolution of the data had a large influence on the accuracy of spatial clustering, regardless of the number of top SVGs selected for the clustering analysis (Fig. S11, Fig. S13). The datasets can be divided into three groups based on their resolutions: a low-resolution group (i.e., resolution > 50μm, such as 10x Visium [50], ST [51, 52]) as shown in Fig. 3C (top) and Fig. 3D, a high-resolution group (i.e., resolution < 50μm, such as Stereo-seq [38] and STARmap [53]) as shown in Fig. 3C (bottom) and Fig. 3D, and a sci-Space group comprising 10 datasets from sci-Space [54]. Although sci-Space can capture barcodes at the single-cell level, it places the coordinates of multiple barcodes at exactly the same position, making it challenging to differentiate between individual cells. Therefore, sci-Space is considered a technology that falls between low-resolution and high-resolution spatially resolved transcriptomic methods, and the 10 datasets from sci-Space were treated as a separate category. The combinations BinSpect+SLM/LVM/Louvain, SOMDE+BayesSpace and SPARK+Louvain performed well on the sci-Space datasets (Fig. S12).

For the low-resolution datasets (Fig. 3C top), the combinations BinSpect+BayesSpace, SPARK+BayesSpace and HSIC+BayesSpace emerged as the best-performing approaches. Spatially aware clustering methods generally achieved higher ARIs compared to scRNA-seq clustering methods (Fig. 3C top), highlighting the importance and necessity of incorporating spatial information for domain assignment even when SVGs are utilized for clustering. Low-resolution datasets often show smoother spatial patterns characterized by gradual transitions in gene expression across the tissue. This inherent smoothness likely contributes to the superior performance of spatially aware clustering methods, since these methods can encourage spatial continuity by leveraging spatial information and gene expression for spatial domain detection.

For high-resolution datasets (Fig. 3C bottom), scRNA-seq clustering methods generally had higher ARIs than spatially aware clustering methods when combined with SVG methods. The combinations of MERINGUE, SpatialDE, SOMDE, BinSpect with three scRNA-seq clustering methods (SLM/ LVM/ Louvain) were the best-performing methods. Among the high-resolution datasets, spatial aware clustering methods only achieved better cluster results for STARmap dataset, possibly due to the simple and smooth region annotations of this dataset (Fig. 3D top, Fig. S14A right). In general, high-resolution spatial technologies can capture finer and more complex spatial structures. Accurate delineation of these intricate spatial domains thus requires spatial clustering methods that can better preserve the granularity and complexity of the observed spatial patterns. Therefore, scRNA-seq clustering methods tend to perform better for high-resolution datasets. Spatial clustering methods with less smoothing enforcement might also work better for high-resolution datasets. However, we found that adjusting the smoothing parameter gamma of BayesSpace to smaller values (gamma=1, 2, 3) had minimal impact on the ARIs of the clustering results (Fig. S15). These findings suggest that further improvement of spatial clustering methods is needed to better handle the complexity of the newer high-resolution spatial data.

For single-cell level spatial transcriptomic data, spatial locations can be clustered to either spatial domains or cell types. To understand the gene selection requirements for these two distinct classification tasks, we conducted clustering analyses for the single-cell level STARmap datasets [53] using cell-type annotations and spatial domain annotations as gold standards, respectively (Fig. S14). In these two experiments, we included top HVG set identified by getTopHVGs [55], which is a state-of-the-art HVG selection method in scRNA-seq data, as an optional input gene set. The results showed that in the combination of SVG/HVG and clustering methods, HSIC+spaGCN performed the best in spatial domain detection tasks, HVG+SLM performed the best in cell-type clustering tasks. In terms of clustering methods, spatial clustering methods were more competent in spatial domain detection tasks due to their consideration of spatial continuity, while scRNA-seq clustering methods that focused on expression profiles were better in cell-type clustering tasks. For SVG/HVG methods in the combination, HSIC, BinSpect, SOMDE and SPARK-X were the best performing methods in spatial domain detection tasks, while MEIRINGUE, HVG, SPARK and SpatialDE outperformed other methods in cell-type clustering tasks. Notably, the combination of scRNA-seq clustering methods and HVG yielded favorable outcomes in cell-type clustering tasks. This observation may indicate that HVG is already competent enough for such cell-type clustering tasks.

### Stability

Adjacent tissue sections or slices commonly display similar gene expression patterns [50]. To evaluate the reproducibility of SVG identification methods, we assessed the consistency of identified SVGs between pairs of adjacent tissue slices. Specifically, we collected 14 pairs of adjacent slices generated using the 10x Visium or ST technologies (Table S1. Sheet3: adjacent slices information). For each method, we calculated the Jaccard similarity between the top 2,000 SVGs identified in each pair of adjacent slices as a measure of reproducibility. As shown in Fig. 4A, SPARK and SpatialDE were the most stable methods, achieving the highest average Jaccard index scores. Furthermore, most methods tended to have higher reproducibility in 10x Visium datasets than ST datasets, which may be due to the fact that the datasets obtained by ST technology only have hundreds of spatial locations and lower spatial resolution. Among them, sepal had the highest reproducibility on the 10x Visium datasets, but its reproducibility levels significantly dropped on the ST datasets, suggesting that the diffusion equation method may be constrained in settings with fewer spatial locations (Fig. S16).

**Fig. 4.**
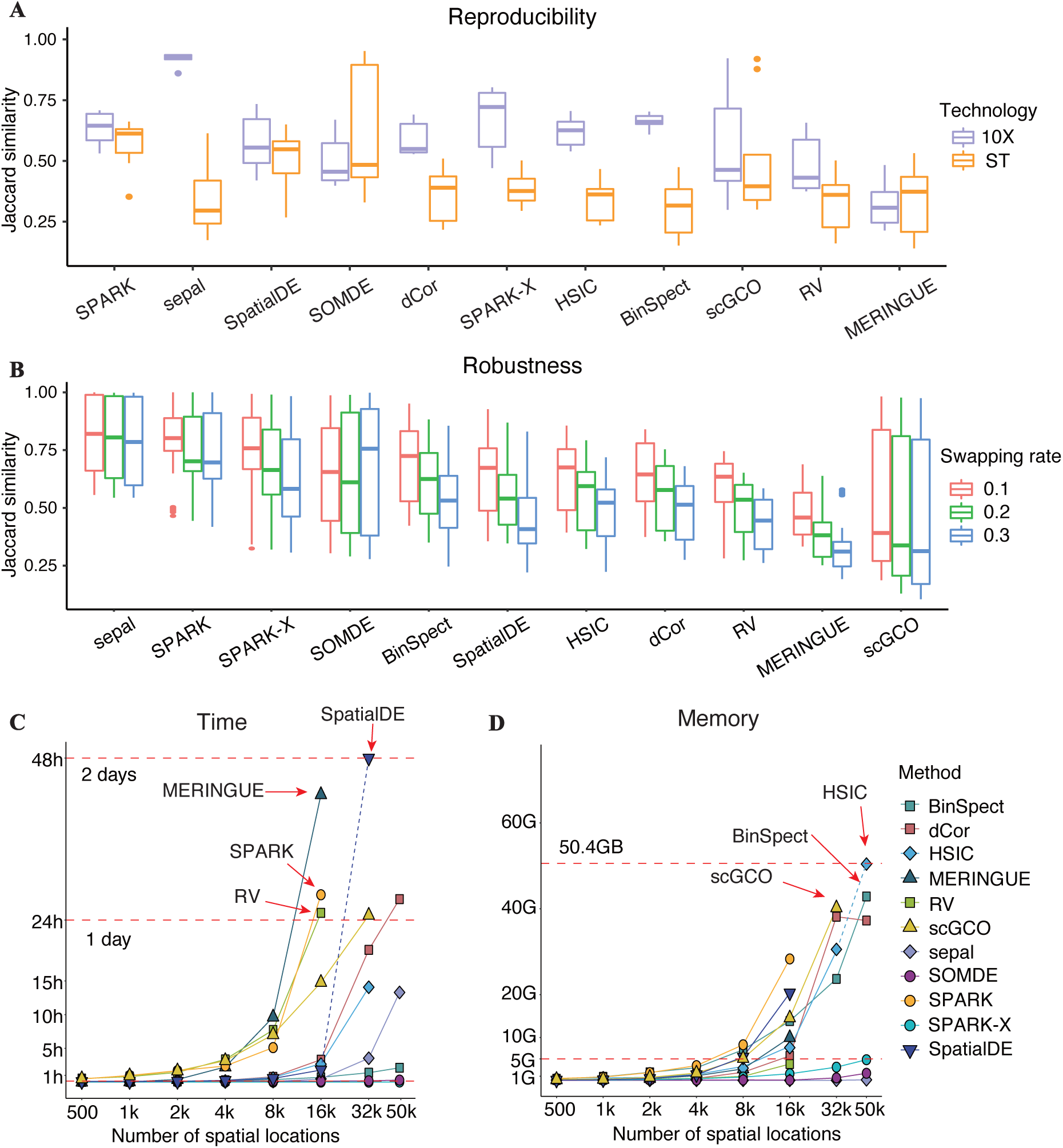
Stability and scalability. **A**, Box plots for the reproducibility measured by Jaccard similarity (the larger, the better) for all adjacent slices, with slices from 10x Visium labeled in purple and slices from ST labeled in orange. **B**, Box plots for the robustness of methods based on Jaccard similarity (the larger, the better), with swapping ratios of *r* = 0.1, 0.2 and 0.3. **C-D**, Scatter plots display the clock time and memory usage of various SVG identification methods for simulated datasets with 10,000 genes and various numbers of spatial locations, with the horizontal axis processed by logarithmic scaling.

Next, we evaluated the robustness of SVG identification methods using perturbed data obtained by randomly “swapping” spatial locations of 11 datasets from various techniques (Table S1. Sheet2: ref. data for robustness). For each gene, we randomly extracted *r*% spatial locations (*r* = 10, 20, 30) and rearranged their expression counts (Methods). This strategy largely preserves the original spatial gene expression patterns (Fig. S17). Then, for each method, the Jaccard index was calculated between the top 2,000 SVGs identified on the perturbed and original datasets. As shown in Fig. 4B and Fig. S18, the Jaccard indices of most methods decreased as the swapping rate increased. sepal, SPARK, and SPARK-X achieved the highest Jaccard indices across all swapping rates. Particularly, sepal demonstrated the strongest robustness across various swapping rates. We also found that as the number of spatial locations increased, most methods demonstrated improved robustness under the location swapping strategy (Fig. S19). This improvement may be attributed to the higher sparsity level of large-scale datasets, where a higher proportion of zero-counts may mitigate the effects of location swapping. Additionally, datasets with more spatial locations may better retain spatial variation in the remaining locations after swapping, further contributing to the robustness of SVG identification methods under such perturbations.

### Scalability

While early SVG methods were developed for analyzing the spatial transcriptomic data with hundreds of spatial locations, current methods need to handle large-scale data with thousands or even tens of thousands of spatial locations. Thus, time cost and computational memory are the key indicators for SVG identification methods to be user-friendly. To evaluate scalability in terms of the time cost and computational memory, we first generated a series of datasets containing 10,000 genes and various numbers of spatial locations (500∼50,000) by down-sampling locations in the Slide-seqV2 dataset[39], and ran the SVG identification methods to each dataset. Each task was executed on a Linux platform with 2.9 GHz Intel Xeon E5-4617 CPUs with 20 parallel threads. The computational time costs and memory usages were recorded. The tasks that exceeded the time limit (48h) and memory limit (50.4 GB) were aborted.

Overall, we found that both computational time cost and memory usages of SVG identification methods increased as the number of spatial locations increased (Fig. 4C-D, Fig. S20). SPARK-X and SOMDE were the fastest methods that successfully completed analyses for all datasets with varying numbers of spatial locations (Fig. S21). BinSpect, sepal finished all tasks within one day. MERINGUE, SPARK, SpatialDE and RV needed more than one day to finish the tasks with more than 16,000 spatial locations. In terms of the computational memory, sepal, SOMDE, and SPARK-X required the least memory usage (less than 5 G for all datasets), while HSIC, scGCO and BinSpect required huge amount of memory (more than 40 GB for datasets with more than 32,000 spatial locations). In summary, considering both computational time cost and memory usage, SOMDE and SPARK-X were the most user-friendly SVG identification methods, particularly for large-scale spatial transcriptomics.

## Discussions

We proposed a comprehensive benchmarking framework designed to evaluate SVG identification methods in spatial transcriptomics. To assess the performance of these methods, we focused on five key criteria. Figure 5 presents a detailed comparison of the characteristics and performance metrics of each algorithm, offering a succinct overview of the benchmarking outcomes. Additionally, it serves as a practical guide, aiding users in choosing the most suitable methods for their specific applications.

**Fig. 5.**
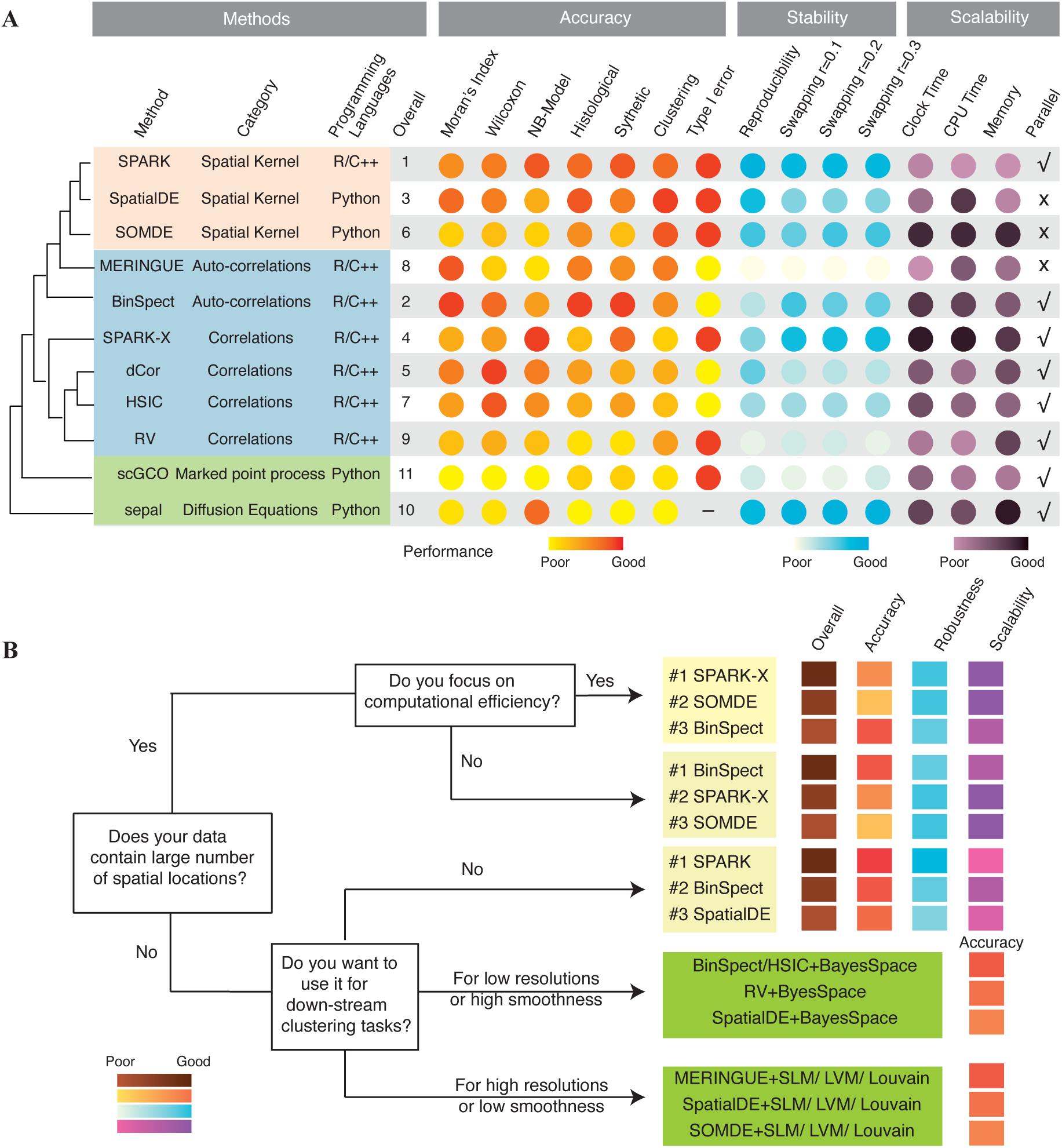
Evaluation summary and recommendation map. **A**, Hierarchical clustering diagram (left panel) displays the similarities between methods. The closer the distance in the tree structure, the higher the similarity of the methods. Comprehensive summary of our evaluations (right panel) including the basic information of the algorithms, their overall ranks, accuracy ranks, robustness ranks and scalability ranks. Each row corresponds to one of the algorithms in our evaluation. Among them, overall ranking is a weighted average of accuracy, robustness and scalability metrics (Methods). **B**, A recommendation map established for specific analysis needs. The bottom left corner graph shows the color representations ranging from poor to good performances. Criteria for performance summary are available in the section Methods.

To facilitate method selection and experimentation by users, we computed the average Jaccard indices for the top 2,000 SVGs identified by different methods (Fig. S21). Using the resulting average Jaccard index profiles, we conducted hierarchical clustering [56] and discovered that the SVG identification methods can be classified into three distinct categories. These categories agreed well with how the spatial information was used by these methods (Fig. 5A, Methods). These methods were implemented in R/C++ or Python with the majority supporting parallelization for better computational efficiency (see Table S2 for details).

Based on the benchmark results, we provided context-specific user recommendations for SVG identification methods (Fig. 5B) as following:

i. Overall, we recommend SPARK, BinSpect, and SpatialDE for their superior accuracies in the majority of datasets.
ii. For large-scale spatial transcriptomic datasets, we recommend SPARK-X and SOMDE for their superior computational efficiencies.
iii. In terms of clustering tasks, users need to assess the resolution, the spatial smoothness of the data, and the specific purpose of clustering. For tasks involving spatial region detection, we recommend choosing a combination of BinSpect/HSIC/RV/SpatialDE and spatial clustering methods (such as BayesSpace) when the data exhibits low resolution or high smoothness characteristics, and a combination of MERINGUE/SpatialDE/SOMDE and scRNA-seq clustering methods (such as Louvain) when the data demonstrates high resolution or low smoothness characteristics. For cell-type clustering tasks, we suggest that a combination of traditional HVG approach and scRNA-seq clustering methods are sufficient.
iv. When users are more concerned with consistency or reproducibility, we recommend SPARK, SPARK-X and BinSpect.

The SVG identification is an active research field, but its evaluation is challenging due to the lack of gold standards. This paper provides a comprehensive evaluation of SVG identification methods and delivers context-specific recommendations for users. As spatial transcriptomic techniques continue to advance towards higher resolution and larger scale, there will be an increasing demand for more robust and efficient identification of SVGs. Based on the benchmark evaluation results, we recommend enhancing the applicability of SVG methods to high-resolution and large-scale datasets. Additionally, we propose integrating SVG methods with downstream analysis tasks, such as spatial clustering and pathway enrichment, to achieve more targeted and informative insights. We anticipate that the benchmark framework will serve as a valuable reference for researchers in developing new SVG identification methods and in applying SVG methods in various spatial transcriptomic studies.

## Methods

### Spatial variable genes identification algorithms

After extensive literature reviews, we include 15 SVG identification methods in our evaluation, comprising 12 publicly available algorithms from the literature and preprints, and 3 additional multivariate-correlation methods (Table S2). A few methods are not included in the evaluation due to the following reasons. SpaGCN [41] and BayesSpace [42] are not included, because these methods primarily focus on clustering analysis and do not specifically develop new methods for SVG identification. The SVG method SpatialDE2 [43] is not included because it lacks the package description and user manual. For all methods, we investigate the authors’ recommendations on how to utilize their methods and follow their guidelines to the best of our ability. For methods that can be executed in parallel, we used 20 CPU cores.

#### Methods based on spatial kernel

For methods using spatial kernels to characterize spatial relations between cells or spots, Gaussian kernel and Gaussian process regression model are mostly applied.

SpatialDE [15] uses a Gaussian process regression model to describe relations between the gene expression and spatial coordinates. In this model, expression variability is decomposed into a spatial variance term and a non-spatial variance term. The spatial variance term is represented by Gaussian kernels related to spatial distance. Genes with a significant spatial variance are reported as SVGs. SpatialDE is widely used and implemented in Python.

SPARK [23] models the gene expression using a quasi-Poisson generalized linear spatial model with a logarithmic link function. To characterize the spatial auto-correlation between spots, SPARK introduces a random effect term with 10 candidate variance kernels, including the Gaussian kernel. Under each candidate kernel, SPARK tests whether a gene shows a spatial expression pattern based on score statistics. Then, SPARK combines the p-values from the 10 kernels using the Cauchy combination rule [57] and adjusts the p-values for multiple-testing with the Benjamini-Hochberg (BH) method in order to obtain a final p-value for each gene.

SOMDE [26] aims to efficiently handle large scale spatial transcriptomic data. It first employs a self-organizing map [58] to aggregate spatially adjacent cells or spots into “nodes”. Then, it identifies node-level SVGs using a Gaussian process approach similar to that used in SpatialDE. By performing spatial aggregation, SOMDE significantly improves computational efficiency.

GPcounts [27] is also based on the Gaussian process regression model. Instead of using a quasi-Poisson generalized linear model as used in SPARK, GPcounts utilizes the negative binomial regression model, because negative binomial distributions have been shown to fit well to bulk RNA-seq data and scRNA-seq UMI (unique molecular identifier) count data [27]. Other components of GPcounts’ model are similar to those of SPARK.

Besides, GPcounts adopts a sparse approximation of variational Bayesian inference to improve computational efficiency. It is implemented using the tensorflow [59] and GPflow [60] in python. However, despite these optimizations, GPcounts still fails to complete computation in 24 hours for more than 90% of the real datasets used in this study.

BoostGP [28] takes the large proportion of zeros in spatial transcriptomics into consideration when setting up the Gaussian process regression model. To account for the potential high dropout rate, BoostGP uses the zero-inflated negative binomial distribution to model gene expression counts. As a Bayesian method, BoostGP uses a MCMC algorithm for Bayesian inference. BoostGP is computationally extensive and fails to complete computation in 24 hours for more than 90% of the real datasets used in this study.

#### Methods based on marked point process

trendsceek [22] employs the marked point process, a technique also prevalent in disciplines such as geostatistics, astronomy and material physics [61], to model the gene expression. Briefly, it models the joint probability distribution of spatial locations and a given gene’s expression. To identify SVGs, trendsceek uses four test statistics: conditional mean (E-mark), conditional variance (V-mark), Stoyan’s mark correlation and the mark-variogram [22]. Its p-values are obtained using random permutation for null model construction, and the accuracy of the p-values depends on the number of permutations. When the number of permutations is 10,000, trendsceek typically requires over 24 hours to complete the analysis for most real datasets used in this study.

scGCO [29] also employs the marked point process to model the gene expression, but it discretizes the noisy expression values into different bins. Hidden Markov random field (HMRF) is used to model the discretized expression values and the latent gene expression states is inferred based on a graph cut algorithm. Then, scGCO tests the dependence between spatial locations’ gene expression states and their spatial coordinates. P-values are derived under the null hypothesis of complete spatial randomness (CSR), or the expression is from the homogeneous spatial Poisson process.

#### Methods based on correlations

One direct approach for detecting SVGs involves assessing correlations between gene expression and spatial coordinates. SPARK-X [30] defines a class of correlations between gene expression and spatial coordinates. To capture various types of expression patterns in real datasets, SPARK-X employs several transformations of the spatial coordinates in calculating the correlations. SPARK-X is computationally efficient even for large-scale sparse datasets.

In addition, inspired by the multivariate correlation perspective [62], we also include three general-purpose correlation coefficients in our benchmark: RV-coefficient [44], distance correlation (dCor) coefficient [45] and Hilbert Schmidt Independent Criterion (HSIC) [46]. One advantage of these general-purpose correlation metrics is that they are model-free. However, they do not incorporate any biological prior knowledge, which thus often necessitates additional screening by users.

Another type of correlations widely used in SVG identification is the so-called spatial auto-correlation [63]. Spatial auto-correlation quantifies the similarity in gene expression between given spots and their nearby spots. The auto-correlation method BinSpect [31] first binarizes gene expression. Fisher’s exact test is then used to determine if the binarized expression exhibits any spatial auto-correlation patterns. BinSpect provides two options for gene expression binarization, by k-means clustering or by sorting and thresholding. To ensure the applicability of BinSpect across datasets, we adopted the k-means method for binarization.

Another auto-correlation method MERINGUE [32] employs Voronoi tessellation to establish preliminary adjacency matrix of spatial locations, and requires adjacent spatial locations to be within a certain spatial distance to confirm their neighborhood adjacency relationships. Leveraging the adjacency matrix and the normalized gene expression as inputs, MERINGUE uses the Moran’s index and local Moran’s index (LISA) for SVG identification.

#### Miscellany methods

BOOST-MI [33] applies a modified Ising model to characterize binarized expression profiles. The network of the Ising model is taken as the nearest neighbor network and the energy interaction parameters represent whether there are spatial patterns in gene expression. The spatial patterns are divided into two types: attraction and repulsion. The repulsion type, a novel spatial pattern proposed in this article, represents a spatial configuration that nearby cells tend to at different expression levels. However, examples of repulsion-type expression in real data are scarce. The algorithm runs for more than 24 hours on most real datasets used in this study.

sepal [34] analogizes the spatial distribution of gene expression to the diffusion process of matter. Taking the observed spatial distribution of gene expression as the initial state, sepal simulates the diffusion process of transcripts through diffusion equations and infers the virtual diffusion time required for each gene to reach a uniform distribution in space. sepal posits the fundamental assumption that genes with random spatial distribution tend to reach uniform distribution states faster than those with structured formation. Consequently, genes with longer virtual diffusion times are considered to have stronger spatial variations. Unlike traditional hypothesis testing frameworks, sepal foregoes p-values, instead relying on virtual diffusion times for ranking SVG.

### Datasets and pre-processing

#### Real datasets

We collect 74 public datasets [38, 50–54, 64–82] from 11 spatial transcriptome technologies, with various spatial structures, resolutions and throughput. The datasets cover a wide range of tissues in humans, mice and other organisms. Detailed information about these datasets can be found in the supplement Table S1.

Among the 74 real-world datasets, 52 have additional information that can aid our evaluation. The datasets with additional information can be divided into two categories. (1) Spatial transcriptomic datasets with region annotations. This category includes 40 public datasets with expert annotations from ST, 10x Visium, sci-Space, seqFISH+, STARmap and Stereo-seq technologies. The expert annotations are treated as the gold standard of region labels; (2) Spatial transcriptomic datasets with aligned histological staining images. This category comprises 12 datasets from 10x Visium technology with H&E images. H&E images contain important histological information and rich evidence of spatial structures [44, 82]. They are commonly used as a main reference by pathologists for manual annotations [52].

#### Synthesized datasets

We generate synthesized data using the SRTsim [40] using three real-world datasets from different technologies as references. Reference dataset 1 is a mouse brain sagittal dataset from the 10x Genomics Visium v1 technology [37], reference dataset 2 is a mouse embryo dataset from the Stereo-seq [38] technology, and reference dataset 3 is a large-scale mouse hippocampal dataset from the Slide-seqV2 [39] technology. The numbers of spots are 2,693, 5,910 and 51,200, for dataset 1, 2 and 3, respectively (Fig. S2). SRTsim requires known domain annotations of the reference dataset for simulating domain specific gene expressions. Expert domain annotations are available for dataset 2, but dataset 1 and 3 lack annotations, so we perform clustering analysis to obtain domain annotations.

For each reference data, we employ SRTsim to systematically choose an appropriate count distribution (one of NB, ZINB, Poisson, or Zero-inflated Poisson) for each gene within every designated region based on expert annotations or clustering outcomes, and then fit the parameters of the selected distribution. With these estimated parameters, we classify all genes to three gene sets, the SVG set, the non-SVG set and the ambiguous gene set. The genes in the ambiguous gene set are those that fall on the boundary of SVGs or non-SVGs. We exclude these genes from the evaluation to prevent potential biases in performance assessment. Let *N* be the number of regions in the reference dataset. Denote *α* = (*α*_1_, *α*_2_, ⋯, *α_N_*) ∈ ℝ*^N^* as the vector of proportions of spatial locations that fall in different annotation regions, and *μ_g_* = (*μ*_1*g*_, *μ*_2*g*_, ⋯, *μ_Ng_*) ∈ ℝ*^N^* as the vector of the estimated mean parameters for gene *g* in each of *N* regions, and *β_g_* = (*β*_1*g*_, *β*_2*g*_, ⋯,*β_Ng_*) ∈ ℝ*^N^* as the vector of the proportions of non-zero counts of gene *g* in each of *N* regions. We define 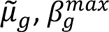 and 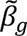 as follows:

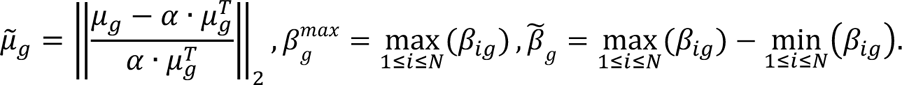

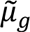 represents the overall deviation of the mean parameters of gene *g* in each of the *N* regions from the grand mean of gene *g*. SVGs generally should have large values of 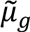 and we can simply define SVGs as those genes with large values of 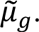 However, a gene can also have a large 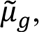 even when the gene does not show overall expression difference between domains but just exhibit abnormally high expression at a small number of isolated points in one domain. To prevent selecting such genes into the SVG set, we require that a SVG must simultaneously have large values of 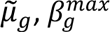 and 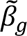 [3]. The thresholds of these three indices are adaptively selected to define the SVG, the non-SVG and the ambiguous gene sets for each reference dataset. Specifically, for reference dataset 1, genes with 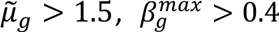 and 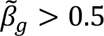 are considered as SVGs, genes with 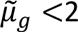 and 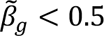 are considered as non-SVGs, and others are considered as ambiguous genes. For reference dataset 2, genes with 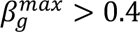 and 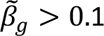 are considered as SVGs, genes 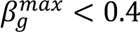 and 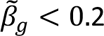 with are considered as non-SVGs, and others are considered as ambiguous genes. For reference 3, genes with 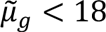 and 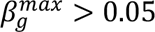 are considered as SVGs, genes with 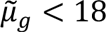 are considered as non-SVGs, and others are considered as ambiguous genes. For each reference dataset, we randomly choose 3000 genes from the SVG set and 7000 genes from the non-SVG set as inputs to SRTsim and generate 10 synthesized datasets.

#### Pre-processing

We apply the same quality control process for all algorithms. For low-resolution datasets (resolution > 20μm), except for GSE111672 [52] dataset from ST technology, we filter out the spatial locations whose library sizes are less than 400 and the genes that are expressed in less than 1% of spatial locations. Specifically, the sequencing depth of GSE111672 datasets appears inadequate, leading to the discard of more than 30% of spatial locations when applying above quality control. To address this issue, we implement a more lenient quality control in this dataset to filter out the spatial locations whose library size less than 200 and the genes expressed in less than 1% spatial locations. For high-resolution datasets (resolution < 20μm), except for GSE130682 [80] datasets from HDST technology, we filter out the spatial locations with library sizes less than 20 and the genes that are expressed in less than 0.1% of spatial locations. The HDST high-resolution datasets are very sparse, so we apply a more lenient quality control. The spatial locations with library sizes less than 5 and the genes that are expressed in less than 5 spatial locations are filtered out. The quality control process aims to retain a broader range of data while still maintaining a reasonable level of quality. The dimensions of the count matrices before and after quality control are shown in the supplement Table S3.

After quality control, the data matrices are then normalized. For the algorithms that specifically developed for SVG detection, we apply their own normalization procedures. For the three general-purpose methods, we use *NormalizeData* from the R package Seurat (v4.0.4) to normalize the data.

### Construction of silver standards

#### Silver standards constructed using spatial aggregation

The Moran’s index is used to test the spatial auto-correlation of gene expression. Then the obtained p-values are adjusted using the Bonferroni method to account for the multiple testing problem. The silver standard genes are selected as those with adjusted p-values less than 0.05. The R package spdep (v1.2_4) is used for calculating the Moran’s indices and the corresponding p-values for spatial transcriptomic datasets with less than 20,000 spatial locations. For large-scale datasets with more than 20,000 locations, the R package moranfast (v1.0) is used for computational efficiency considerations.

#### Silver standards constructed using the Wilcoxon test

Let *N* be the number of regions in the expert annotations. For each gene, the Wilcoxon rank-sum test is used to test whether there is a significant difference in expression levels between any pair of regions. This test is performed for *N*(*N* − *1*)/*2* possible pairs of regions. The *N*(*N* − *1*)/*2* p-values by comparing pairs of regions are then combined as a single p-value using the Cauchy method [57]. Then, the combined p-values for all genes are adjusted using the Bonferroni method to account for the multiple testing problem. Genes with adjusted p-values less than 0.05 are taken as the silver standard SVGs. The Wilcoxon test is performed using the R package stats (*wilcox.test*) v4.0.5, and the Cauchy method for combing p-values is implemented using the *ACAT* function in R package ICSKAT.

#### Silver standards constructed using the NB regression

Given the expert annotation, let *X*_*i*_ be the vector of dummy variables corresponding to the annotated region labels of spatial location *i*, and *N*_*i*_ be the library size of spatial location *i*. Given gene *g*, denote *Y_gi_* as its expression counts in spatial location *i*, and 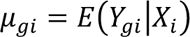 as its conditional mean. We consider the following negative binomial generalized linear model

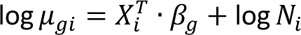

where *β_g_* is the vector of regression coefficients. When *β_g_* = *0*, gene *g* is considered to be a non-SVG. We use the likelihood ratio test to test the hypothesis *β_g_* = *0*. The p-values for all genes are using the Bonferroni method to account for the multiple testing problem. Genes with adjusted p-values less than 0.05 are taken as the silver standard SVGs. The NB regression and the likelihood ratio tests are performed using R packages MASS (v7.3_54) and stats (v4.0.5).

#### Silver standards constructed by correlating with histology images

After converting the RGB image data to lightness vectors through the LAB [83, 84] transform, we test each gene’s expression correlation with the image lightness using Pearson’s correlation. As before, genes with Bonferroni corrected p-values less than 0.05 are selected as SVGs.

#### Accuracy metrics calculation

Utilizing the silver standard SVGs in real datasets and genuine SVGs in synthetic datasets, we rank genes according to each SVG method and then compute three widely recognized accuracy metrics: AUPR, AUROC, and EP. AUPR and AUROC are calculated by function *pr.curve* and *roc.curve* in R package PRROC (v1.3.1). EP is calculated as the fraction of true positives in the top-K identified SVGs, where K is the number of the sliver SVGs. The higher values for these three accuracy metrics indicate better performance of the SVG detection method.

### Clustering analysis

The identified SVGs are used as features for clustering analysis. We apply scRNA-seq clustering methods Louvain algorithm [47], LVM [48], and SLM [48], as well as the spatial-aware clustering methods BayesSpace [42] and SpaGCN [41] to cluster the spatial locations. Given the expert annotations, we tune the parameters of these five clustering methods, including the “resolution” parameter of the scRNA-seq clustering methods, the “q” parameter of BayesSpace, the “target_num” parameter of SpacGCN, such that the resulting number of clusters equals to the number of clusters in the expert annotations.

### Computation of the statistical validity criteria

For each method except for sepal, the detected SVGs from a spatial transcriptomic dataset are taken as the genes for which the null hypotheses are rejected at a nominal FDR level of 0.05. In other words, the genes whose BH adjusted p-values are less than 0.05 are considered as the detected SVGs. Since sepal does not provide statistical p-values, we follow the methodology outlined in the sepal paper and apply the function *processStream* from the R package cpm (v2.3) to find a suitable inflection point that serves as the truncation point for the scaled average diffusion times curve. For sepal, we define the SVGs as the genes whose scaled average diffusion times greater than this inflection point. The proportion of SVGs is calculated as the ratio between the number of detected SVGs and the total number of genes remained after pre-processing.

To estimate the type I errors of SVG detection algorithms, for each of the 11 real datasets (Table S1. Sheet2: ref. data for robustness), we construct pseudo datasets by randomly permuting each gene’s expression values across the spatial locations. We then apply the SVG methods to the pseudo datasets. The proportions of genes identified as SVGs by each method at a nominal p-value cutoff of 0.05 are served as an empirical estimate of the type I error rates. Since sepal does not provide p-values, it is not considered for the type I error analysis.

### Computation of the stability criteria

The reproducibility of a method is calculated as the Jaccard similarity between the top-2,000 SVGs detected in adjacent slices of spatial transcriptomics. The reproducibility is calculated for 14 pairs of adjacent slices from 10x Visium and ST technologies (the supplement Table S1. Sheet3: adjacent slices information).

To evaluate SVG methods’ robustness against the “spot-swapping” perturbation, we first generate perturbed datasets for each of 11 datasets from various technologies (Table S1. Sheet2: ref. data for robustness). Given a real dataset, for each gene, we randomly select *r*% spatial locations (*r* = 10, 20 or 30), and then randomly shuffle the gene’s expression values on the selected spatial locations. We repeat this process 10 times to generate 10 perturbed datasets with a swapping rate of *r*% for the given real dataset. We then apply each SVG method to the perturbed datasets and calculate the Jaccard similarity between the top-K SVGs detected in the perturbed datasets and the original real dataset. We take *K* = 2,000 for all datasets except the STARmapMBR [53] datasets. The total number of genes in STARmapMBR is only 1020, so we use *K* = 200. This swapping strategy largely preserves the raw spatial patterns in gene expression (Fig. S17).

### Computation of the scalability criteria

To test the scalability of SVG methods, we construct datasets of various sizes by randomly down-sampling spots of Slide-seqV2 dataset [39]. The numbers of spatial locations of the down-sampled datasets range from 500 to 50,000. Each SVG method is applied to the down-sampled datasets on a Linux platform with 2.9 GHz Intel Xeon E5-4617 CPUs with 20 parallel threads. The clock time, CPU time and memory usage are recorded for each method on each down-sampled dataset. We set a 48-hour clock time limit and a 50.4-GB memory usage limit. Tasks exceeding these limits are aborted. The memory usage is recorded using the *peakRAM* function in the R package peakRAM (v1.0.2) or the *Profile* function in the python profile (v0.61.0) package.

### Hierarchical clustering of SVG methods

For each of the 74 real datasets, we calculate Jaccard similarity for the top-K genes for each pair of methods. We take *K* = 2,000 for all datasets except the STARmapMBR [53] datasets. The total number of genes in STARmapMBR is only 1020, so we use *K* = 200. Then, the hierarchical clustering [56] is applied to cluster the methods based on the Jaccard similarity matrix. The function *hclust* in the stat R package is used for the hierarchical clustering.

### Evaluation pipeline

#### Input

Real or synthesized spatial transcriptomic datasets after pre-processing are used as inputs.

#### Software

For the algorithms that specifically developed for SVG detection, the softwares we used are BinSpect (Giotto v1.0.4), SPARK (SPARK v1.1), MERINGUE (MERINGUE v1.0), SpatialDE (SpatialDE v1.1.3), SOMDE (somde v0.1.8), SPARK-X (SPARK v1.1.1), sepal (sepal v1.0.0), scGCO (scGCO v1.1.0), trendsceek (trendsceek v1.0.0), BOOST-GP (Github documentation download required), BOOST-MI (Github documentation download required), GPcounts (GPcounts v0.1). For the three general-purpose correlation methods, we use R packages FactoMineR (v2.4), Rfast (v2.0.6) and dHSIC (v2.1) to calculate the correlations RV, dCor and HSIC between spatial locations’ coordinates and gene expression. Detailed information about these algorithms can be found in the supplement Table S2.

#### Parameter selection

For all SVG methods in our benchmark, we investigate recommendations provided by the authors and strive to implement these suggestions to the best of our ability. Most parameters are set as suggested by the user manuals of the SVG algorithms. For the few parameters that require user specification or are not set as their default values, we set them as described below.

1. For MERINGUE, the certain distance parameter is used to filter neighborhood adjacency relationships constructed by Voronoi tessellation. Drawing from the author’s diverse illustrations utilizing real-world data, the distance parameter is set as either 2, 2.5, or 10, contingent upon the specific characteristics of the data. To enhance MERINGUE’s capability for generalization across multiple datasets, we refrain from explicitly specifying this distance parameter.
2. For SOMDE, the parameter that determines the degree of aggregation of spatial locations needs user specification. We employ *SomNode(coordinate, 20)* for datasets with more than 20,000 spatial locations, *SomNode(coordinate, 10)* for datasets with 2,000-20,000 spatial locations, and *SomNode(coordinate, 4)* for datasets with less than 2,000 spatial locations.
3. For trendsceek, we set its permutation parameter to 10,000 for more precise p-value estimation.

#### Output

The outcomes of each SVG identification algorithm encompass the genes’ ranks and their associated p-values or scores given by the algorithm, as well as the runtime and memory usage of the algorithm. For all algorithms except sepal, the genes are ranked based on the adjusted p-values given by the algorithm. In the case of sepal, since it does not perform statistical testing and does not provide p-values, the gene ranks are calculated using sepal’s gene selection criterion—the scaled average diffusion times.

For each SVG algorithm, we evaluate SVG detection accuracy, statistical validity, accuracy of downstream clustering, stability and scalability in accordance with established standards, and output the evaluation results.

### Performance summary of SVG methods

We rank the SVG methods using each of the evaluation metrics. Given a SVG method, let 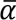 be its average rank of accuracy metrics, 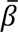 be its average rank of robustness, and 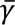 be its average rank of scalability. We define the overall rank score as 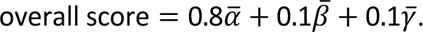

For large scale datasets, when users are more concerned about computational efficiency, we recommend the algorithms based on the rank score 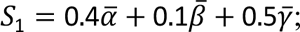 When users are less concerned with computational efficiency, we recommend the algorithms based on the rank score 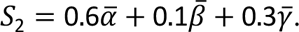

If clustering analysis is the major goal, we make recommendations based on the average ARI rankings of the 55 combinations of methods across all datasets with expert annotation information; Otherwise, we make recommendations based on the overall rank scores.

## Data availability

All real datasets considered in this study are publicly available and can be download from Zenodo (https://doi.org/10.5281/zenodo.7227772). Details about these datasets are provided in the supplement Table S1.

## Code availability

A R/Python implementation of the benchmark framework is available at https://github.com/XiDsLab/svg-benchmark.

## Acknowledgments

This work was supported by the National Key R&D Program of China [2020YFE0204200 to R.X.], the National Natural Science Foundation of China [11471022, 71532001 to R.X.], and Sino-Russian Mathematics Center. Part of the analysis was performed on the high-performance computing platform of the Center for Life Sciences (Peking University).

## Author Contributions

Conceptualization, R.X.; Methodology, J.T., Z.C. and X.C.; Software: X.C., Q.R.; Formal Analysis: Z.C., J.T., X.C. and Q.R.; Resources: X.C. and S.H.; Writing, X.C., R.X. and X.S.; Funding Acquisition, R.X.; Supervision, R.X. and X.S.

